# Adding iPSC donor lines does not adequately control for genetic heterogeneity

**DOI:** 10.64898/2026.04.22.720258

**Authors:** Artur Shvetcov, Shannon Thomson, Caitlin A. Finney

## Abstract

Human induced pluripotent stem cell (iPSC)-based disease modelling studies are widely expected to include three to five independent donor lines to control for the contribution of donor genetic background to phenotypic variance. This convention has been formalized into major guidelines, yet no power analysis has evaluated whether these sample sizes can detect, estimate, or control for donor-level genetic effects. Here, we provide that evaluation. Using Monte Carlo simulation, closed-form confidence intervals, population genetics, and empirical resampling of transcriptomic data from iPSC lines, we show that studies with three to five donors cannot reliably detect donor-level variance, cannot estimate its magnitude with useful precision, and cannot determine whether a treatment effect generalizes across genetic backgrounds. The sample sizes required to reliably detect, estimate, or control for donor-level variance exceed 20 donors and, for many phenotypes, exceed 50, well beyond what any standard disease modelling experiment can deliver. Adding two or three donor lines to a study does not meaningfully increase statistical power, narrow confidence intervals, or establish whether a treatment effect generalizes across genetic backgrounds. The inability to control for genetic background is not a limitation of individual study design but a structural property of iPSC-based modelling. We propose that the field adopt isogenic controls for variant-specific questions and orthogonal validation against clinical datasets for generalizability, rather than treating donor number as a proxy for rigour.

## Introduction

Human induced pluripotent stem cells (iPSCs) are somatic cells, typically dermal fibroblasts, peripheral blood mononuclear cells, or renal epithelial cells collected from urine, reprogrammed to a pluripotent state through forced expression of Yamanaka factors ^1,2^.

Unlike human embryonic cells, iPSCs can be derived from virtually any adult donor, preserving the donor’s genomic background and circumventing the ethical constraints associated with embryonic sources. Once established, iPSC lines can be expanded indefinitely and directed toward differentiation into many somatic lineages through combinations of morphogens, small molecules, and genetic manipulations ^3^. Over the past decade, this capacity has transformed *in vitro* modelling of human development and disease, progressing from two-dimensional monolayer cultures into three-dimensional organoid systems that recapitulate tissue architecture, cellular diversity, and functional physiology ^4-6^. Brain intestinal, liver, kidney, lung, and cardiac organoids provide platforms for studying organ-specific disease mechanisms ^6-9^ and assembloids combining multiple regional identities enable the study of inter-tissue interactions previously accessible only *in vivo* ^10-12^.

This methodological expansion has coincided with a rapid shift in the regulatory environment. The FDA Modernization Act 2.0 eliminated the statutory requirement for animal testing in preclinical drug development and explicitly recognized iPSC-derived models, organoids, and organ-on-chip systems as acceptable alternatives ^13^. The FDA’s 2025 roadmap articulated a trajectory in which animal studies become the exception rather than the norm for certain therapeutic classes within three to five years. Here, human cell-based new approach methodologies will serve as primary evidence for safety and efficacy evaluation ^14^. Publications using iPSC-derived models have accelerated markedly in parallel ^7,15^, placing a corresponding burden on the field to establish experimental standards that can bear the weight of translational decision making. Chief among the standards in current use is the requirement that iPSC-derived studies include multiple donor lines to address the well-documented contribution of donor genetic background to phenotypic variance. Consortium-scale studies have estimated this contribution at between 5 and 46% ^16^. Recommendations have been made to use three, four or five donor lines per experimental group, with the logic that sampling multiple genetic backgrounds controls for their contribution to the observed phenotype ^17-21^. These recommendations have since been formalized into journal editorial policies, with ISSCR guidelines encouraging journals to require data from multiple patients and controls ^18,20,21^. However, neither the original recommendations nor the editorial policies derived from them were accompanied by power analyses demonstrating that these sample sizes achieve meaningful sensitivity to detect, estimate, or control for donor-level genetic effects.

In this article, we present statistical, genetic, and empirical evidence that this logic does not hold at the donor counts feasible for disease modelling studies, providing the missing quantitative evaluation of these widely adopted thresholds. A study with one donor and a study with five donors contains effectively the same amount of information about donor-level genetic effects, almost none. The sample sizes required to meaningfully detect or generalize across donor-level variance, on the order of more than 20, are beyond the reach of any standard disease-modelling experiment, and no incremental increase from three to five or five to ten changes this. The inability to control for donor genetic background is not a deficiency of experimental design. It is a structural property of iPSC-based disease modelling that cannot be engineered away. We argue that the field should stop treating donor number as a proxy for rigor, acknowledge this limitation explicitly, and invest instead in strategies that genuinely strengthen inference. These include isogenic controls for variant-specific questions, orthogonal validation against human tissue data, and experimental designs matched to questions the system can answer.

### Characterizing genetic heterogeneity in iPSCs requires population-scale studies

The quantitative characterisation of donor-level variance in iPSC systems has been achieved not through routine individual laboratory studies but through dedicated consortium-scale efforts. Early work established that genetic background, rather than reprogramming method or cell type of origin, is the dominant driver of transcriptional variation between iPSC lines ^22^. The Human Induced Pluripotent Stem Cells Initiative (HipSci) remains the most comprehensive of these, characterising 711 iPSC lines from 301 healthy donors through standardized reprogramming and multi-modal phenotyping ^16^. Donor genetic background explained between 5 and 46% of observed phenotypic variance, consistently exceeding contributions from sex, passage number, reprogramming method, and culture conditions. The resulting iPSC expression quantitative trait locus (eQTL) map identified thousands of regulatory variants active in the pluripotent state ^16^. Complementary efforts have extended these findings across independent cohorts and modalities. The NHLBI NextGen Consortium profiled 317 iPSC lines from 101 donors and attributed approximately 50% of genome-wide expression variability to inter-individual differences ^23,24^. Matched proteomic and transcriptomic analysis of 202 HipSci lines from 151 donors demonstrated that donor-level variance persists at the protein level even after accounting for transcript abundance, implicating genetically driven post-transcriptional regulation that is invisible to RNA profiling alone ^25^. Single-cell RNA-sequencing from 125 HipSci donors during directed endoderm differentiation further revealed that genetic effects on expression are dynamic rather than static, with hundreds of eQTLs emerging, disappearing, or changing direction across differentiation stages ^26^. Similarly, profiling of iPSC-derived sensory neurons from HipSci donors demonstrated that donor genetic effects on gene expression persist after differentiation into a mature somatic cell type, confirming that genetic background is not diluted by the differentiation process ^27^. Together, these studies establish genetic background as the single largest identifiable source of molecular and cellular variation in iPSC systems.

### Why population-scale approaches cannot be universalized

A logical corollary of this consensus might appear to be that all iPSC studies should adopt population-scale designs. Such designs are feasible and informative when the research question is fundamentally epidemiological, quantifying how a phenotype varies across healthy genetic backgrounds or mapping the genetic architecture underlying that variation. However, several constraints preclude their generalization to the much broader category of iPSC studies concerned with disease mechanisms, therapeutic targets, or cellular pathophysiology.

Recognizing the potential for batch and culture-associated variance in large-scale parallel differentiation, researchers have adopted pooled village designs in which iPSCs from multiple donors are differentiated in a single dish and de-multiplexed *post hoc* using donor-specific single nucleotide polymorphisms ^26,28-31^. This design inherently allows for large numbers of iPSC donors and batch effects are eliminated as a confound as every donor line is exposed to identical conditions. However, pooled village designs are well suited only to studying variation among biologically similar donors under shared conditions and cannot be extended to most disease modelling contexts. When iPSC lines from healthy controls and disease affected individuals are co-cultured, cells interact through paracrine and autocrine signalling, compete for nutrients, and modulate one another’s differentiation trajectories.

Even when whole genome sequencing is performed on donor lines, it is used for demultiplexing and genetic association mapping rather than systematic screening for deleterious variants that could affect the shared culture environment. Ostensibly healthy donors may therefore carry rare pathogenic or functionally consequential variants whose cellular effects propagate through the village and confound the phenotypes of neighbouring lines. Pooling such lines would therefore introduce a novel artifact in which each donor’s phenotype is shaped by its neighbours. For example, iPSC-derived neurons, astrocytes, and microglia-like cells carrying the apolipoprotein E ε4 variant (*APOE* ε4) secrete distinct profiles of cytokines and lipoproteins that would alter neighbouring cell physiology ^32^. In a pooled village format, every iPSC line would be exposed to this disease-modified microenvironment, confounding any comparison between genotypes as well as disease cases versus controls. Even among nominally healthy lines, iPSC line-specific competitive dynamics have been observed, with some lines lost from the pool within days of initiating differentiation ^28^. Consistent with this, reanalysis of a pooled dopaminergic neuron differentiation experiment across 238 iPSC lines revealed that individual lines become overrepresented by up to 3-to 10-fold within a single pool ^30,31^. Somatic mutations in developmental genes such as *BCOR* were strongly associated with both differentiation failure and elevated proliferation rates, suggesting that competitive dynamics in village formats are driven in part by genetic variation ^30^. Any disease state that alters proliferation, survival, or differentiation efficiency, which includes most neurodegenerative, oncological, and developmental disorders, would systematically distort donor representation and invalidate the intended comparison(s).

These constraints leave most disease modelling laboratories in the position that population-scale and pooled village designs are both inaccessible. The prevailing recommendation, to include three to five independently derived donor lines per experimental group has emerged as a practical compromise ^17-19^. The implicit assumption is that while three to five donors cannot capture the full spectrum of genetic variation, they provide meaningful control for donor background effects. The analyses that follow test this assumption directly.

### Small donor designs are statistically uninformative for detecting genetic heterogeneity

To quantify the information about donor-level genetic effects contained in studies of typical size, we performed Monte Carlo simulations of hierarchical variance component estimation. Donor counts ranged from 2 to 50, and true intraclass correlation coefficient (ICC; the proportion of total phenotypic variance attributable to differences between donors) values spanned 5% to 50%, reflecting the empirically observed range from HipSci ^16^. For each simulation, six replicate measurements per donor were modelled under a one-way random-effects ANOVA, and power, ICC point estimates, confidence intervals, and Bayesian posterior probabilities were computed (Supplemental Information).

The simulation results expose a fundamental limitation. At a true ICC of 20%, roughly the median of the HipSci range, a study with three donors achieves only 26% power to detect the donor effect. Five donors raise this to 37% and 10 donors to 58%, all below the conventional 80% threshold. Power reaches 82% only at 20 donors (Figure 1a,b). For phenotypes at the lower end of the empirical range (ICC = 5%), such as cell morphology traits where donor identity explains as little as 8% of variance ^16^, power at n = 3 is 9.5%, barely distinguishable from the 5% false positive rate. At these sample sizes, the statistical test cannot differentiate between the presence and absence of a donor-level genetic contribution.

**Figure 1.**
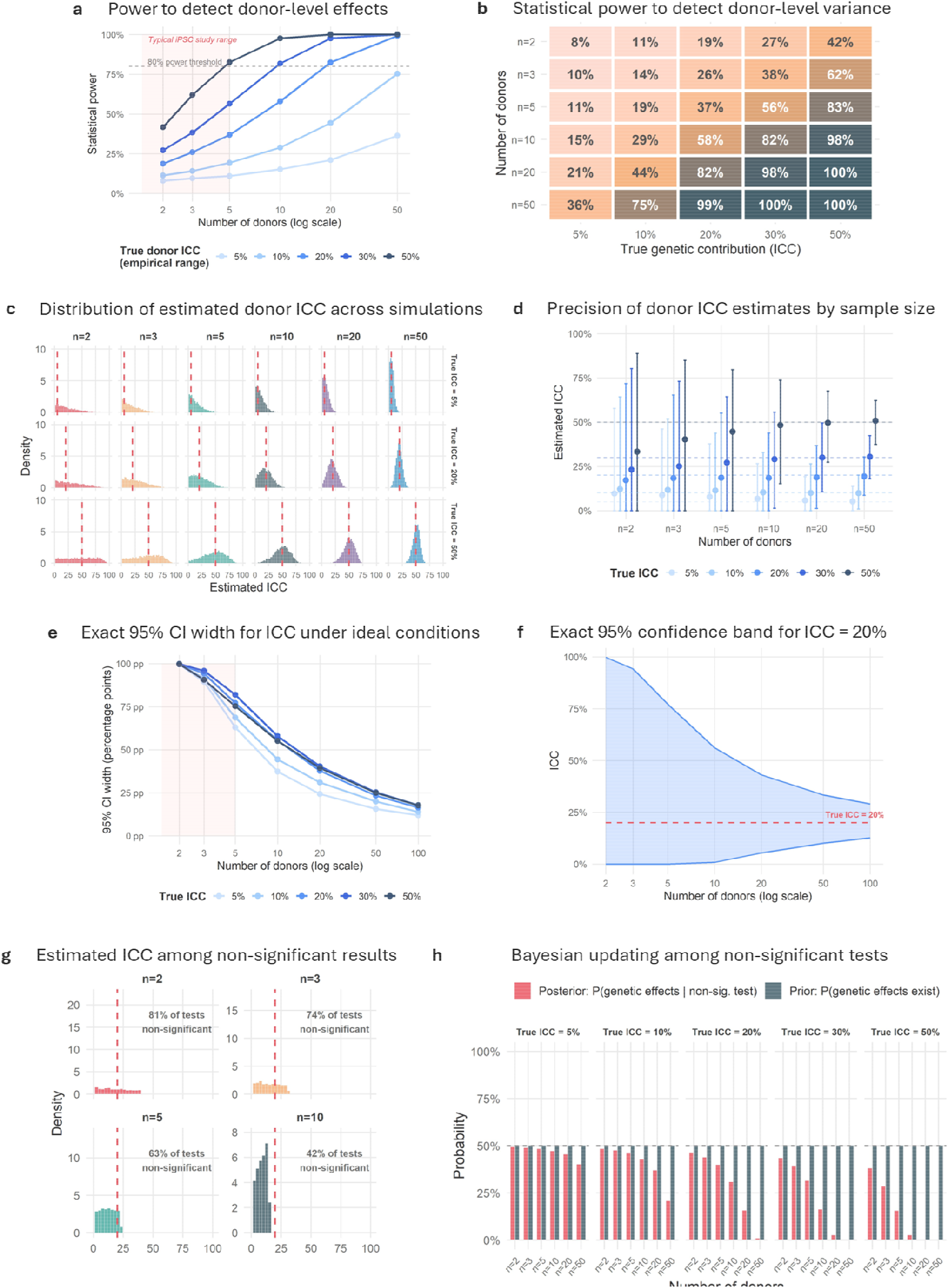
Small donor designs cannot detect or estimate donor-level genetic effects. (**a**) Power curves showing the probability of rejecting the null hypothesis of no donor variance as a function of the number of donors (n = 2 to 50), for five values of the true interclass correlation coefficient (ICC = 5%, 10%, 20%, 30%, 50%). Each point represents the proportion of 5,000 Monte Carlo iterations yielding p < 0.05 with k = 6 replicates per donor. The horizontal dashed line marks 80% power. (**b**) Heatmap of rejection probability over a grid of donor counts and true ICC values, with the 80% power contour overlaid. The region corresponding to typical iPSC study designs (n = 2 to 10, ICC = 5% to 30%) is almost entirely below 50% power. (**c**) Distribution of ICC point estimates across 5,000 simulated studies for selected donor counts and true ICC values, illustrating that at n = 2 to 5, estimates span the nearly full parameter space regardless of the true value. (**d**) Mean ICC estimate with 2.5^th^ to 97.5^th^ percentile ranges across 5,000 simulated studies at each combination of donor count and true ICC. Dashed horizontal lines mark the true values. At n = 5 or fewer, the empirical 95% range spans most of the parameter space. (**e**) Expected width of the exact 95% confidence interval for donor ICC as a function of donor count, computed from the F-distribution under balanced one-way random-effects ANOVA with k = 6 replicates per donor. At n = 3, expected CI widths exceed 89 percentage points for every tested ICC. A CI width below 20 percentage points requires approximately 100 donors at ICC = 20%. (**f**) Expected exact 95% confidence band for ICC at a true value of 20%. At n = 5 or fewer, the band spans from 0% to above 90%, encompassing the full range of qualitative conclusions. (**g**) Distribution of ICC point estimates restricted to simulations in which the donor F-test was non-significant (p > 0.05), at a true ICC of 20%. At n = 2, 81% of tests are non-significant, and the resulting estimates span the full interval, demonstrating that a non-significant donor test at these sample sizes carries no information about the true ICC. (**h**) Bayesian posterior probability that a donor effect exists given a non-significant F-test, computed from the power values in (**a**) under a uniform prior of 0.5. At realistic ICC values and typical sample sizes, a non-significant result shifts the posterior minimally from the prior.

The problem extends beyond power to the precision of the estimates themselves (Figure 1c,d). At two donors with a true ICC of 20%, the 95% simulation interval for the estimated ICC spans 0% to 72%; at three donors, 0% to 65%. An estimate of 0% and an estimate of 60% are equally routine outcomes from the same underlying biology. This imprecision is compounded by systematic bias at the zero boundary. When between-donor variance falls below within-donor variance by sampling chance, the estimator is truncated at zero. This occurs in 32% of simulations at three donors and true ICC = 20%, rising to 52% at true ICC = 5%. These zeros do not reflect an absence of donor effects but rather the inability of the design to resolve them. Closed-form confidence intervals from the F-distribution, which depend on no simulation assumptions, reinforce this conclusion. At three donors with six replicates and a true ICC of 20%, the expected 95% CI spans 0% to 94% (Figure 1e,f). A single study at this sample size cannot distinguish between a system in which donor identity is irrelevant and one in which it dominates. Achieving a CI width below 20 percentage points, a reasonable minimum for scientifically interpretable precision, requires approximately 100 donors at ICC = 20%.

These findings also reframe the interpretation of non-significant donor tests. When a study with two to five donors reports no significant effect of donor identity, this is frequently taken as evidence that genetic background does not meaningfully contribute to the phenotype.

At two donors and true ICC = 20%, 81% of tests fail to reach significance, yet the underlying donor contribution remains 20% throughout (Figure 1g). Even increasing to five donors still leaves 63% non-significant results. A Bayesian analysis makes this explicit. starting from an uninformative prior of 0.5, a non-significant test at three donors and true ICC = 20% shifts the posterior probability that donor effects exist from 50% to 44% (Figure 1h). The posterior is virtually unchanged from the prior. Therefore, a reported absence of a significant donor effect at these sample sizes reflects the limitations of the test, not the absence of the effect.

### Three to five donors do not sample the genetic landscape

Independent of any statistical consideration, a population genetics analysis reveals that small donor panels fail to capture the genetic variation they are intended to represent. To quantify this, we computed the probability that at least one copy of a variant at a given minor allele frequency is present among 2n sampled haplotypes, given by 1-(1-p)^2n^, where p is the minor allele frequency and n is the number of diploid donors. This was integrated over defined minor allele frequency (MAF) ranges, common (5-50%), low frequency (1-5%), and rare (0.1-1%), under a neutral folded site frequency spectrum, which assumes that allele frequencies follow the distribution expected in the absence of selection. For a biallelic variant with a MAF of 5%, three donors (six haplotypes) have a 74% probability of not carrying even a single copy; at MAF = 10%, the probability of complete absence is 53% (Figure 2). Integrated across the common variant spectrum (MAF 5% to 50%), three donors sample approximately 64% of available variation, leaving more than a third unobserved (Figure 2).

**Figure 2.**
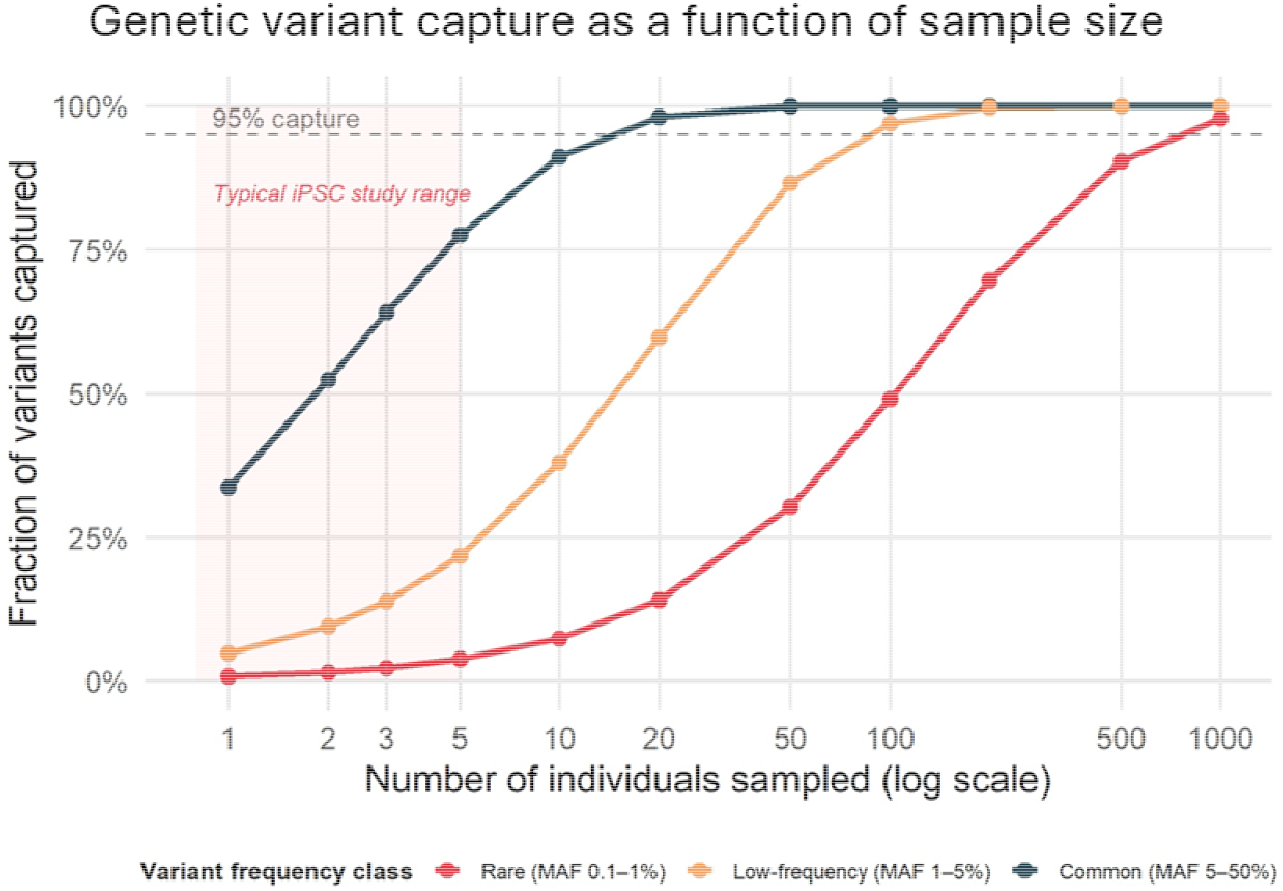
Three to five donors capture a small fraction of human genetic variation. Fraction of variants present in at least one of 2n sampled haplotypes, computed as 1-(1-p)^2n^, integrated over the specified minor allele frequency (MAF) range under a neutral folded site frequency spectrum. Three curves correspond to common (MAF 5% to 50%), low-frequency (MAF 1% to 5%), and rare (MAF 0.1% to 1%) variants. The shaded band marks the typical iPSC study sample size range (n = 1 to 5) and the dashed line marks 95% capture. At n = 3, sampling captures 64% of common variants, 14% of low-frequency variants, and 2% of rare variants. Increasing to n = 5 provides only marginal improvement (78%, 22%, and 4%, respectively). Reaching 95% capture requires approximately 20 donors for common variants, 100 for low-frequency variants, and more than 1,000 for rare variants.

Coverage is far worse for low frequency variants (MAF 1% to 5%), where only 14% are captured, and for rare variants (MAF 0.1% to 1%), where the figure falls below 3% (Figure 2). Increasing the sample to five donors, the upper bound of most current recommendations, provides only modest improvement covering 78% of common variants, 22% of low-frequency variants, and 4% of rare variants (Figure 2). The marginal gain from three to five donors is small relative to the scale of what remains unsampled. Meaningful representation of low frequency variation requires tens of donors, and rare variation requires hundreds. For reference, the HipSci cohort of 301 donors ^16^ captures greater than 99.9% of common variants and approximately 97% of low-frequency variants. This is the scale of sampling required to make credible claims about genetic contributions to phenotypic variance. It is also incompatible with iPSC-derived disease modelling, where generating and differentiating even 10 donor lines in parallel for numerous assays represents a substantial undertaking.

### Empirical resampling confirms the theoretical predictions

To validate the simulation findings empirically, we used published transcriptomic data from Mirauta et al. (2020), comprising batch-corrected, log-transformed expression values for 9,013 protein-coding genes across 202 iPSC lines from 151 donors ^25^. We computed a reference set of donor ICCs for each gene from the 51 donors with two or more iPSC lines (Supplemental Information). The median donor ICC across all genes is 43.2%; 90% of genes exceed 20%, and 35% exceed 50% (Figure 3a). At this scale, donor identity accounts for more expression variance than any other measured factor, consistent with earlier reports from HipSci ^16^ and the NHLBI NextGen Consortium ^23^.

**Figure 3.**
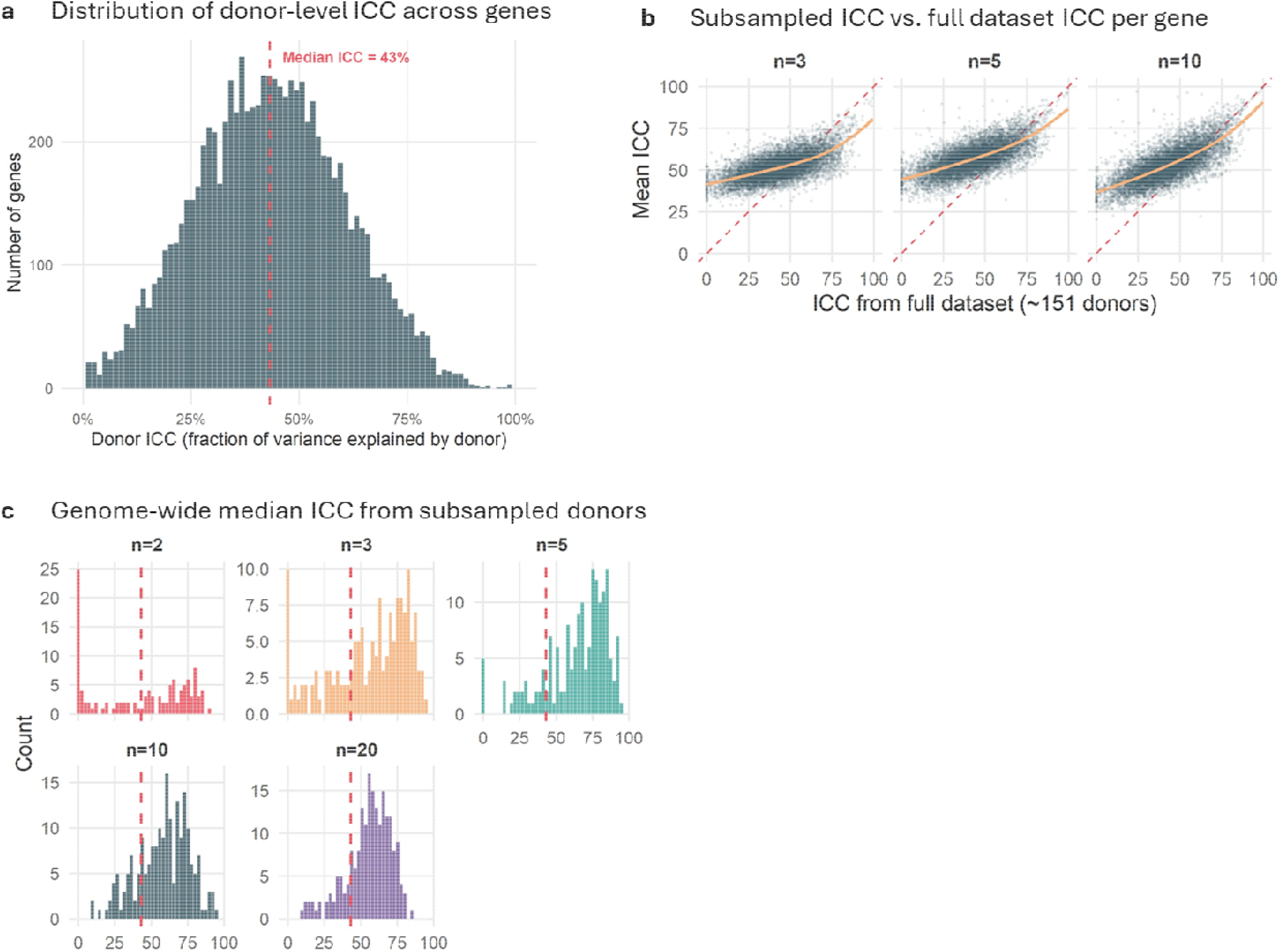
Empirical resampling of large-scale transcriptomic data confirms that small-donor estimates are unreliable. (**a**) Distribution of gene-wise donor ICCs across 9,013 protein-coding genes, computed by one-way random-effects ANOVA from the 51 donors with two or more iPSC lines in the Mirauta et al. (2020) dataset ^25^. The red dashed line marks the median donor ICC (43.2%). Over 90% of genes have ICC above 20% and 35% exceed 50%. (**b**) For each of 9,013 genes, the mean donor ICC across 200 random donor subsamples (y-axis) plotted against the ICC from the 51-donor reference (x-axis), for subsamples of n = 3, 5, 10, and 50 donors. The red dashed line shows identity, and the orange curve is a loess smooth. At n = 3, subsampled estimates compress toward the middle of the distribution with a fitted slope of approximately 0.36, and the per-iteration 95% range spans 0% to 99.8%. (**c**) Distribution of the genome-wide median donor ICC across 200 random subsamples of 2, 3, 5, 10, 20, and 50 donors. The red dashed line marks the reference median (43.2%). At n = 3, the median subsampled estimate is 62.1% with a 95% range of 0% to 91%, demonstrating that the reported genome-wide donor contribution at typical iPSC study sample sizes is driven as much by which donors were sampled as by the underlying biology.

We then subsampled donors at sizes typical of published studies (n = 2 to 50) and re-estimated gene-wise ICCs (Supplemental Information). At three donors, the relationship between subsampled estimates and the reference is severely attenuated, with a fitted slope of 0.36 between reference ICC values of 10% and 90%. Subsampled estimates compress toward the middle of the distribution regardless of the true value. A gene with a reference ICC of 10% and a gene with a reference ICC of 70% return overlapping estimate distributions. If a three donor subsample returns an ICC estimate of 60%, the true value is plausibly anywhere between 10% and 76% (Figure 3b). The point cloud tracks the diagonal only at 20 donors or above, and at two donors, 42% of random subsamples cannot be analyzed at all because every sampled donor contributes only a single iPSC line. The simulation predicted that at three donors, ICC estimates would span 0% to 65% regardless of the true value. The empirical resampling confirms and exceeds this, with per-gene 95% ranges spanning 0% to 99.8% at a reference ICC of 10%. The empirical noise meets or exceeds what the simulation predicted under idealized conditions. This instability of small sample estimates also extends beyond individual genes to genome-wide summary statistics. When the genome-wide median donor ICC was computed for each random subsample, the distribution at three donors was centered at 62.1%, with a 95% range spanning 0% to 91%, compared to the 51-donor reference median of 43.2% (Figure 3c). At these sample sizes, the reported genome-wide contribution of donor identity is driven as much by which donors happened to be sampled as by the underlying biology.

### Small donor panels cannot determine whether an effect generalizes across genetic backgrounds

A related but distinct question is whether the effect of a treatment or perturbation iPSC experiment is consistent across genetic backgrounds, rather than driven by a subset of donors. Statistically, this asks whether the donor-by-treatment interaction variance differs from zero. To evaluate this, we simulated a balanced two-way mixed design across a range of donor counts and interaction magnitudes (Supplemental Information).

When every donor responds identically to treatment, five donors achieve 82% power to detect the mean effect at an effect size of one residual standard deviation, broadly consistent with existing sample size recommendations ^17,18,20^ (Figure 4a). However, these recommendations assume homogeneity of response across donors. Once the treatment effect is permitted to vary, power deteriorates rapidly. At a donor-by-treatment interaction standard deviation of 0.7, where individual donors deviate from the mean treatment response by amounts comparable to the within-donor noise, the same mean effect yields only 32% power at five donors and 15% at three. This occurs because the correct denominator for testing the mean treatment effect is the interaction mean square, not the residual. When donors respond differently to treatment, this denominator grows and the test loses sensitivity. Sample size recommendations derived from iPSC models that assume uniform response across donors will therefore underestimate the number of donors required.

**Figure 4.**
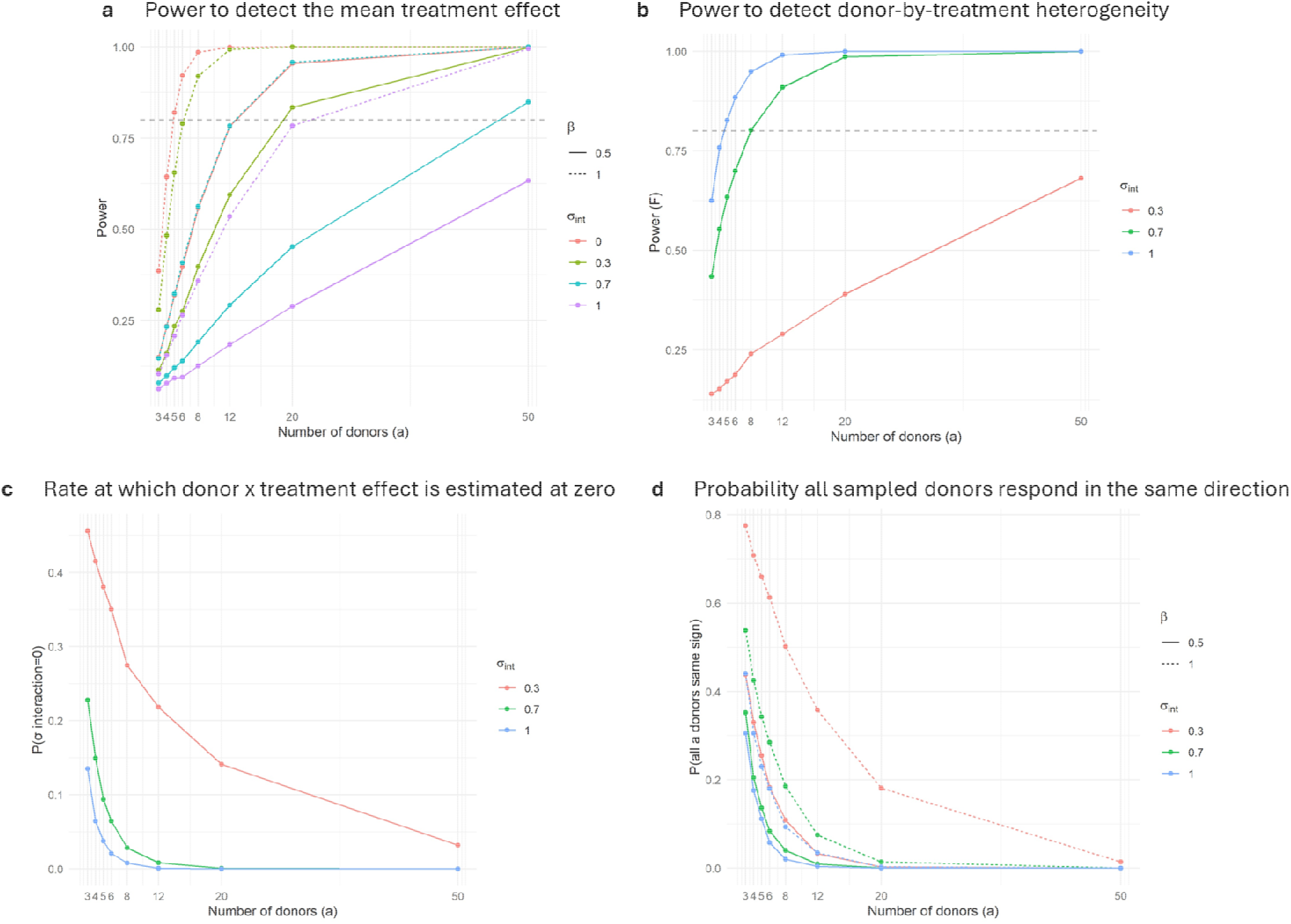
Small donor panels cannot determine whether treatment effects generalize across genetic backgrounds. (**a**) Power curves for the correctly specified F-test of the mean treatment effect across donor counts (n = 3 to 50), for combinations of treatment effect size and donor-by-treatment interaction standard deviation. In the homogeneous response case, five donors achieve 82% power for a treatment effect of one residual standard deviation. Under realistic heterogeneity (interaction SD = 0.3 to 0.7), the same mean effect yields 15% to 32% power at n = 5. (**b**) Power curves for the F-test of the interaction variance component across donor counts and interaction magnitudes. At moderate heterogeneity (interaction SD = 0.3), power does not reach 80% at any tested sample size up to 50. (**c**) Proportion of simulations in which the method-of-moments estimate of the interaction variance is truncated to zero, across donor counts and true interaction standard deviations. At interaction SD = 0.3, 47% of three donor iterations and 33% of 6-donor iterations return zero estimates, not because heterogeneity is absent but because the design cannot resolve it. (**d**) Probability that all sampled donors show treatment effects of the same sign, across donor counts and true interaction standard deviations. At interaction SD = 0.3 and a treatment effect of one residual standard deviation, 77% of three donor studies and 66% of five donor studies appear unanimous across donors. The visual impression of consistency at small sample sizes is produced primarily by the number of donors sampled rather than by genuine uniformity of response.

Testing whether the effect generalizes across donors, rather than simply whether it exists on average, is even more demanding. To evaluate this, we simulated a two-way mixed design in which multiple donors each receive a treatment and a control condition, and the treatment effect is allowed to vary across donors. The degree of this variation is captured by the donor-by-treatment interaction standard deviation (SD), a measure of how much individual donors deviate from the average treatment response. When this value is zero, every donor responds identically; as it increases, donors diverge in how strongly, or even in which direction, they respond. At moderate heterogeneity (interaction SD = 0.3, meaning individual donors deviate from the mean response by roughly a third of the within-donor noise), the interaction F-test achieves 13% power at three donors and 18% at five, and does not reach 80% at any sample size up to 50 (Figure 4b). The variance estimator is truncated at zero in 47% of three donor iterations, not because heterogeneity is absent but because the design cannot resolve it (Figure 4c).

The same problem applies to visual assessment of consistency. Researchers often judge whether a treatment effect is robust by checking whether all donors respond in the same direction (e.g. all donors show increased cell death following treatment). At an interaction SD of 0.3 and a treatment effect of one residual standard deviation, 77% of three donor studies and 66% of five donor studies will show all donors responding in the same direction (Figure 4d), despite meaningful underlying heterogeneity in response magnitude. At these parameter values, a three donor study has 28% power to detect the mean effect, 14% power to detect response heterogeneity, a 46% chance of estimating the interaction variance at exactly zero, and a 77% chance of appearing consistent across all donors. Such a study would typically be reported as demonstrating a robust effect replicated across three independent genetic backgrounds. That conclusion, however, is not supported by the underlying data.

These numbers place the required sample sizes far beyond what iPSC-derived disease modelling can realistically deliver. A study with upwards of 20 donor lines would require each line to be independently reprogrammed, validated for pluripotency and karyotypic stability, and differentiated through protocols that often span weeks to months per line. Every downstream assay, whether proteomic, transcriptomic, electrophysiological, or imaging-based, would need to be performed across all lines with appropriate biological and technical replication. This rapidly multiplies reagent costs, instrument time, and analytical complexity by an order of magnitude. For rarer diseases, the constraint is even more fundamental. The patient populations themselves may not contain enough individuals willing and able to donate or perhaps patients are spread out globally, making the required sample sizes both impractical and impossible. Even for common diseases, no single laboratory or consortium has demonstrated the capacity to run a fully replicated complex iPSC-derived modelling experiment across this many donors for the range of assays typical of a mechanistic study.

### Isogenic controls can help answer variant-specific questions

If small donor designs cannot adequately control for genetic background effects, what experimental strategies remain? When the objective is to determine whether a specific genetic variant produces a cellular phenotype, the isogenic comparison is the most rigorous available design. Here, a patient-derived iPSC line is paired with a variant corrected isogenic control or a healthy donor line is engineered to carry the variant of interest, such that the resulting pair differs exclusively at the target locus ^33,34^. Phenotypic differences observed in this framework can be attributed to the variant with a degree of confidence that no number of unmatched donors can achieve. Adding unmatched donors to an isogenic experiment does not strengthen inference about the variant, it introduces uncontrolled genomic variation that obscures the comparison of interest. A related argument holds that multiple donor lines serve as a practical screen for line-specific artifacts such as karyotypic abnormalities or failed differentiations.

However, this conflates quality control with genetic background control. Gross line-specific failures are more reliably detected through systematic characterization at line establishment than through informal comparison across a handful of unmatched donors ^20,35,36^. A sample of three to five provides no statistical basis for distinguishing a line-specific artifact from normal inter-donor variation. The isogenic design can specifically answer whether a given variant produces a measurable phenotype on the genetic background under study. It does not, however, establish whether the same variant produces the same phenotype across all backgrounds. As the analyses above demonstrate, this broader question is not answerable at any donor count feasible for standard disease modelling experiments. If the donor counts required for meaningful inference about genetic background effects cannot be achieved, then the inability to control for this variable is not a limitation that will be resolved by future investment or scale. It is an inherent property of iPSC-based disease modelling. A study with three donors and a study with one donor contains equivalent information about donor-level genetic contributions, and treating these two designs as qualitatively different misrepresents the statistical foundation on which both rest.

### Orthogonal validation provides what donor replication cannot

If neither small donor replication nor the infeasible alternative of large donor replication can establish that an iPSC-derived finding generalizes across genetic backgrounds, the question becomes what can. We argue that the most productive strategy is orthogonal validation against human biological data from clinical cohorts. When findings from an iPSC-derived model are independently recapitulated in clinical cohort data, whether from tissue, biofluids, or large-scale genomic studies, the convergence across platforms and sample types provides stronger evidence of biological relevance than adding further iPSC donor lines. Although this approach remains underutilized, there are studies that demonstrate its value. For example, iPSC-derived cardiomyocytes recapitulated the clinical susceptibility of individual breast cancer patients to doxorubicin-induced cardiotoxicity ^37^. In Alzheimer’s disease, Wang et al. (2025) integrated large-scale proteomic and genetic data from post-mortem brain to build causal network models of disease, then validated key predictions in iPSC-derived astrocytes from a single donor ^38^. Taking a complementary approach, our team compared proteomic profiles from iPSC-derived cortical organoids against peripheral and central immune profiles from a clinical cohort of *APOE* ε4 carriers, identifying convergent dysregulation biological pathways ^39^. In each case, confidence in the findings derives from agreement between the iPSC-derived model and independent human data, not from the number of donor lines used. We suggest that such comparisons against clinical datasets should become routine, as they address the generalizability concern far more directly than incremental increases in donor number.

### Recommendations for the field

Based on these findings, we propose a revised framework for how iPSC-derived disease modelling studies should be designed, interpreted, and reviewed.

1. Studies should be designed around questions the experimental system can answer. When the objective is to determine whether a genetic variant produces a cellular phenotype, the isogenic comparison remains the gold standard, and the experiment should be evaluated on those terms. When the objective is to assess whether a treatment rescues a disease phenotype, the comparison should be made within an isogenic or single donor framework with appropriate biological replication, rather than diluted across unmatched donors whose genomic differences introduce variance that the study is not powered to resolve.
2. Claims about genetic heterogeneity should be commensurate with the data. Studies should not state or imply that they have controlled for, accounted for, or addressed donor genetic background unless they have statistically shown this. We have shown here that this is unlikely to be achievable. The number of donors used should be reported as a design parameter, and the inability to generalize across genetic backgrounds should be stated as an inherent constraint of the iPSC-derived modelling rather than treated as a shortcoming of the individual study.
3. Orthogonal validation should replace donor replication as the primary strategy for establishing generalizability. Demonstrating that iPSC-derived findings converge with independent clinical datasets, whether from tissue, patient biofluids, or population-scale genomic studies, provides direct evidence of human relevance that no feasible increase in donor number can match.
4. Reviewers and editors should reassess the practice of requesting additional donor lines as a condition of publication. Adding two or three donors to a study does not address genetic heterogeneity in any statistically meaningful way and risks creating an illusion of rigor where none exists. The relevant question is not how many donors were used but whether the study is designed to answer a question that the iPSC-based system is equipped to address. The ISSCR Standards for Human Stem Cell Use in Research ^20^ provide a valuable framework for promoting rigor and reproducibility and the quantitative analyses presented here may help inform future iterations of these recommendations as the field continues to refine best practices for experimental design.
5. Where population-level questions are genuinely central to the study, pooled village approaches should be adopted in contexts where non-cell-autonomous interactions between genotypes do not confound the results. For disease modelling contexts where pooling is not feasible, population-level questions about genetic architecture should be pursued through complementary approaches such as GWAS, eQTL mapping, or clinical cohort studies, and not forced into an experimental system that lacks the statistical power to resolve them.

## Conclusions

The practice of including three to five donor lines in iPSC-derived disease modelling studies as a means of controlling for donor genetic heterogeneity is not supported by statistical theory, Monte Carlo simulation, population genetics, or empirical reanalysis of large-scale transcriptomic data. Across every analysis presented here, the conclusion is the same, a study with one donor and a study with five donors contain effectively the same amount of information about donor-level genetic effects. These designs differ in appearance, not in statistical rigor or their ability to control for differences in donor genetic backgrounds. Despite this, iPSC-derived models remain among the most powerful experimental systems available for studying human disease. They provide access to living human cell types that cannot be obtained by any other means, they enable mechanistic dissection of disease-relevant pathways in the cells affected by disease, and they are increasingly central to preclinical therapeutic development. None of this is diminished by the findings presented here. What is diminished is the notion that adding a handful of donor lines transforms a single-background observation into a generalizable one.

The field faces a choice. It can continue to treat donor number as a proxy for rigor, requiring three to five lines per study despite the absence of statistical justification for this threshold. Or it can acknowledge that genetic background dependence is a structural property of iPSC-based disease modelling and report this limitation transparently. The strategies that do strengthen inference are already available. Isogenic designs address variant-specific questions, orthogonal validation against human clinical data addresses generalizability, and experimental designs matched to answerable questions address credibility. Rigor does not require pretending a limitation has been solved, it requires designing around it.

## Author contributions

A.S.: Data curation, formal analysis, methodology, software, visualization

A.S. and C.A.F.: Conceptualization, funding acquisition, project administration, resources, supervision, Writing – Review & editing

A.S., S.T., and C.A.F.: Investigation, writing – original draft

## Acknowledgements

This work was supported by funding from the Australian Government’s National Health and Medical Research Council Medical Research Future Fund MRF2040081 (C.A.F. and A.S.) and MRF2052401 (A.S. and C.A.F.); philanthropic funding from the John & Anne Leece Family (A.S.), Paul & Valeria Ainsworth Family (C.A.F.) and Neil and Norma Hill Foundation (C.A.F.).

## Data and code availability

The methods used in the current article are described in detail in Supplemental Information. This study used publicly available data as described in Supplemental Information. Code and simulation outputs are available at https://github.com/Art83/donor_iPSC.

