## Supplementary Information for "Adding iPSC donor lines does not adequately control for genetic heterogeneity"

**Supplemental Information**

**Empirical grounding of simulation parameters**

Intraclass correlation coefficient (ICC) values used in the simulations were selected to span the empirically observed range reported by consortium-scale studies. The donor-level ICC is defined as the proportion of total phenotypic variance attributable to differences between donors. An ICC of 0 means that knowing which donor a measurement came from tells you nothing about the measurement’s value; all observed variation is replicate-level noise. An ICC of 1 means every replicate from a given donor is identical, and all variation in the dataset comes from differences between donors. In practice, the ICC for a given phenotype falls somewhere between these extremes and is defined as:

ICC = *σ^2^_donor_* / (*σ^2^_donor_* + *σ^2^_residual_*)

where *σ^2^_donor_* is the variance attributable to differences between donors (how much donors differ from one another on average) and *σ^2^_residual_* is the residual variance (how much replicate measurements from the same donor scatter around that donor’s mean). When *σ^2^_donor_* is large relative to *σ^2^_residual_*, the ICC is high and donor identity strongly shapes the phenotype. When it is small, the ICC is low and replicate noise dominates.

We selected ICC values of 0.05, 0.10, 0.20, 0.30, and 0.50 to span the empirically observed range. Kilpinen et al. (2017) characterised 711 iPSC lines from 301 donors and reported that inter-individual differences explained 5-46% of phenotypic variance depending on the assay. Donor identity was the dominant source of gene expression variance for 46.4% of microarray probes and cell morphological features showing donor contributions of up to 23% ^1^. Carcamo-Orive et al. (2017) profiled 317 iPSC lines from 101 individuals and attributed approximately 50% of genome-wide expression variability to inter-individual differences ^2^. Mirauta et al. (2020) performed matched quantitative proteomics and RNA-sequencing on 202 HipSci iPSC lines from 151 donors and demonstrated broadly distributed donor-level variance at the protein level ^3^. Our five ICC values therefore cover the full range from phenotypes barely influenced by donor identity (5%) through those moderately shaped by it (20-30%) to those dominated by it (50%).

**Monte Carlo simulation of donor-level variance estimation**

The goal of this simulation was to determine what happens when a researcher runs an iPSC experiment with a small number of donors and then tries to assess how much of the observed variation is due to donor identity. To do this, we modelled a balanced one-way random-effects design in which each of *a* donors contributes *k* = 6 replicate measurements. For each simulation iteration, we generated data according to a known ground truth. Donor-level effects were sampled from a pool of 1,000 virtual donor values drawn from a normal distribution with mean zero and standard deviation (SD) *σ_donor_*_._ This SD was back calculated from each target ICC value as:

*σ_donor_* = √(ICC x *σ^2^_residual_* / (1 - ICC))

Within-donor residuals, representing the scatter of replicate measurements around each donor’s true mean, were drawn from a normal distribution with mean zero and *σ_residual_* = 1. By fixing the residual variance at 1 and varying only the donor variance, we could systematically control how much of the total variation was due to donor identity (the ICC) while keeping the simulation interpretable. Donor sample sizes of *a* (2, 3, 5, 10, 20, and 50) were evaluated across all five ICC values, with 5,000 iterations per condition.

For each simulated dataset, we performed a one-way analysis of variance (ANOVA), which partitions the total variation into two components: variation between donor group means (captured by the between-group mean square, MS_between_) and variation among replicates within donors (captured by the within-group mean square, MS_within_). The between-group mean square reflects both the true differences between donors and the replicate noise, while the within-group mean square reflects replicate noise alone. When donors truly differ, MS_between_ will tend to be larger than MS_within_; the ratio of these two quantities forms the F-statistic used to test the null hypothesis that there is no between-donor variance. The ICC was then estimated from these mean squares using the standard formula:

ICC = (MS_between_ - MS_within_) / (MS_between_ + (k - 1) x MS_within_)

This estimator works by subtracting the replicate noise (MS_within_) from the total between-group variation (MS_between_) to isolate the donor contribution, then expressing it as a proportion of the total. However, when the number of donors is small, sampling fluctuations can produce a situation where MS_between_ falls below MS_within_ purely by chance, even though donors truly differ. In these cases, the formula returns a negative value, which is biologically meaningless, so the estimate is truncated at zero ^4^. This truncation does not mean that donors are identical; it means the experiment lacked the resolution to see their differences.

For each condition, we recorded the proportion of iterations achieving p < 0.05 (statistical power), the mean and median ICC estimates, the 2.5^th^ - 97.5^th^ percentile simulation intervals, and the proportion of iterations yielding an ICC estimate of exactly zero.

**Exact confidence intervals from the F-distribution**

In addition to the Monte Carlo simulations, we computed confidence intervals for the ICC using an exact analytical method that does not rely on simulation. For a balanced one-way random-effects model with a donors and k replicates per donor, the expected F-ratio (the ratio of between-donor to within-donor variability) relates to the ICC by:

*F* = 1 + *k* x ICC / (1 - ICC)

which can be rearranged to express the ICC in terms of the F-ratio:

ICC = (*F* - 1) / (*F* + *k* - 1)

Following Thomas and Hultquist (1978) ^5^, exact 95% confidence bounds were obtained by computing lower and upper F-limits:

*F*_L_ = *F* / *F*_0.975, df1, df2_	and	*F*_U_ = *F* / *F*_0.025, df1, df2_

and then transforming each through the inverse relationship above to yield confidence bounds on the ICC, truncated to the interval [0, 1]. Here, df1 = *a* - 1 and df2 = *a*(*k* - 1) are the degrees of freedom from the ANOVA. The lower F-limit uses the 97.5^th^ percentile critical value (dividing by a large number produces a small bound), and the upper uses the 2.5^th^ percentile (dividing by a small number produces a large bound), together capturing 95% of the sampling distribution.

These intervals were evaluated at the expected population F-value for each combination of *a* donor sample sizes (2, 3, 5, 10, 20, and 50), *k* = 6, and each of the five ICC values. This approach provides a best-case scenario. It represents the expected confidence interval width under ideal conditions of perfect balance, exact normality, no missing data, and no batch confounding. Any departure from these assumptions in real experiments would produce wider intervals.

**Bayesian posterior update from a non-significant donor test**

When a study with a small number of donors reports a non-significant F-test for the donor effect, this is often interpreted as evidence that donor genetic background does not meaningfully contribute to the phenotype. To evaluate how much information such a result actually carries, we computed the Bayesian posterior probability that donor effects exist, given a non-significant test result. Starting from an uninformative prior that assigns equal probability to two hypotheses: (1) *H*_1_: donor-level genetic effects exist, with *P*(*H*_1_) = 0.5; and H_0_: they do not, with *P*(*H*_0_) = 0.5. The posterior was computed via Bayes’ rule:

*P*(*H*_1_ | non-sig) =

*P*(non-sig | *H*_1_) × P(*H*_1_) / [P(non-sig | *H*_1_) × P(*H*_1_) + P(non-sig | *H*_0_) × P(*H*_0_)]

*P*(non-sig | *H*_1_) = 1 – power is the probability of getting a non-significant result when donor effects truly exist (obtained from the Monte Carlo simulations described above), and *P*(non-sig | *H*_0_) = 0.95 is the probability of getting a non-significant result when donor effects are truly absent (i.e. 1 - the false positive rate). Because statistical power at small donor counts is so low, *P*(non-sig | *H*_1_) is close to *P*(non-sig | *H*_0_), meaning a non-significant test is nearly as likely when donor effects exist as when they do not. The posterior therefore barely moves from the prior, confirming that a non-significant donor test at these sample sizes provides almost no evidence either way.

**Population genetics: Variant representation by sample size**

Independent of the statistical power considerations above, we asked a more fundamental question: does a panel of three to five donors even contain a representative sample of human genetic variation? For a biallelic variant with minor allele frequency (MAF) *p*, the probability that at least one copy appears among n randomly sampled diploid individuals (each carrying two copies of the genome) is:

*P*_capture_ = 1 – (1 - *p*)^2^*^n^*

This formula gives the probability that a variant at frequency *p* in the population will be present in a sample of *n* donors. The exponent 2*n* reflects the fact that each diploid donor contributes two haplotypes (one from each parent), so *n* donors provide 2*n* opportunities to observe the variant.

To compute the expected fraction of all variants in a given frequency class that would be captured, this probability was integrated over the site frequency spectrum (the distribution of allele frequencies across the genome). We used a neutral folded site frequency spectrum approximation, which assumes allele frequencies follow the distribution expected in the absence of natural selection, with density proportional to 1/*p*. Under this model, rare variants are far more numerous than common ones, making them the dominant class of human genetic variation and the most poorly sampled. Three variant frequency classes were evaluated: common (MAF 5-50%), low frequency (MAF 1-5%), and rare (MAF 0.1-1%).

**Empirical resampling of HipSco transcriptomic data**

To test whether the simulation predictions hold in real data, we used published transcriptomic data from Mirauta et al. (2020), comprising processed, batch-corrected, log-transformed gene-level expression values for 9,013 protein-coding genes across 202 iPSC lines from 151 donors ^3^. A reference set of gene-wise donor ICCs was computed from the 51 donors who had two or more iPSC lines (102 lines total) using the same one-way random-effects ANOVA estimator described above, with donor as the grouping factor. This subset represents the only portion of the data in which within-donor (clone-to-clone) variance is directly observable, which is required for computing the ICC.

To simulate what would happen if smaller studies were drawn from the same population, we subsampled donors at target sizes of n = 2, 3, 5, 10, 20, or 50. For each target size and each of 200 independent random draws, *n* donors were sampled without replacement from the full set of 151, and gene-wise ICCs were re-estimated. Draws in which all sampled donors contributed only a single iPSC line were recorded as inestimable rather than contributing zeros, because the ICC cannot be computed without within-donor replication. Gene-level misclassification was assessed using practically meaningful thresholds. Here, a gene with a reference ICC above 20% that received a subsampled estimate below 10% was classified as a false negative (a real donor effect missed by the small study), and a gene with a reference ICC below 10% that received a subsampled estimate above 20% was classified as a false positive (a spurious donor effect created by sampling noise).

**Donor-by-treatment interaction simulation**

The simulations above address whether small studies can detect or estimate donor-level variance. A separate question is whether they can determine if a treatment effect is consistent across donors. In other words, whether all donors respond similarly to a perturbation, or whether some respond strongly, some weakly, and some perhaps in the opposite direction. Statistically, this is captured by the donor-by-treatment interaction. We simulated a balanced two-way mixed design with *a* donors x 2 conditions (control, treated) x *k* = 6 replicates per cell. The data-generating model was:

*y_dck_* = *α_c_* + *u_d_* + *v_dc_* + *ε_dck_*

Each term in this model represents a distinct source of variation. The measurement *y_dck_* for the *k*^th^ replicate of donor d under condition *c* is the sum of: *α_c_*, the fixed effect of condition (with *α_control_* = 0 and *α_treated_* = β, the treatment effect size); *u_d_*, the random effect of donor *d* (how far that donor’s baseline deviates from the overall mean); *v_dc_*, the donor-by-treatment interaction (how much donor *d*’s treatment response deviates from the average treatment response); and *ε_dck_*, the residual noise for that particular replicate.

The interaction term *v_dc_* is the quantity of primary interest in this simulation. When its standard deviation (*σ_int_*) is zero, every donor responds identically to treatment. As *σ_int_* increases, donors begin to diverge in how they respond. Some may show a large treatment effect, others a small one, and at high values some may even respond in the opposite direction. We varied *σ_int_* across 0, 0.3, 0.7, 1.0, spanning the full range from perfectly uniform response through heterogeneity comparable in magnitude to the within-donor noise. The donor variance *σ_donor_* and residual variance *σ_residual_* were both fixed at 1, while the treatment effect size β was varied across 0.5, 1.0. Donor counts ranged from 3 to 50 with 5,000 iterations per condition.

Three F-tests were computed from the expected mean squares of the unrestricted mixed model. For the fixed treatment effect, the test statistic is the ratio of the treatment mean square (MS_C_) to the interaction mean square (MS_DC_), with 1 and *a* - 1 degrees of freedom. This test uses the interaction mean square, rather than the residual, as its denominator because when donors respond differently to treatment, this heterogeneity inflates the apparent treatment effect and must be accounted for. For the interaction itself, the test statistic is MS_DC_ / MS_E_ with *a* - 1 and 2*a*(*k* - 1) degrees of freedom, where MS_E_ is the residual mean square. For the donor main effect, the test statistic is MS_D_ / MS_DC_ with *a* - 1 and *a* - 1 degrees of freedom. Variance components were recovered by the method of moments and truncated at zero when negative. Null calibration was verified by 20,000 iterations with *β* = 0 and *σ_int_* = 0, confirming empirical type I error rates near the nominal 0.05.

Four outcomes were recorded per condition: (Q1) power to detect the mean treatment effect; (Q2) power to detect the donor-by-treatment interaction; (Q3) the rate at which the method-of-moments estimator of the interaction variance was truncated to zero; and (Q4) the probability that all a sampled donors displayed a treatment effect of the same sign. This last outcome quantifies how often a small-donor study will appear visually consistent across donors when the underlying treatment response is genuinely heterogeneous.

**Software**

All simulations and analyses were implemented in R (version 4.5.1). No iterative mixed-model fitting procedures were employed; the ANOVA F-tests and method-of-moments variance component estimators are closed form and exact under balanced designs. Code and simulation outputs are available at <https://github.com/Art83/donor_iPSC>.
